# SpatialView: An interactive web application for visualization of multiple samples in spatial transcriptomics experiments

**DOI:** 10.1101/2023.06.13.544836

**Authors:** Chitrasen Mohanty, Aman Prasad, Lingxin Cheng, Lisa M. Arkin, Bridget E. Shields, Beth Drolet, Christina Kendziorski

## Abstract

Spatial transcriptomics (ST) experiments provide spatially localized measurements of genome-wide gene expression allowing for an unprecedented opportunity to investigate cellular heterogeneity and organization within a tissue. Statistical and computational frameworks exist that implement robust methods for pre-processing and analyzing data in ST experiments. However, the lack of an interactive suite of tools for visualizing ST data and results currently limits the full potential of ST experiments. To fill this gap, we developed SpatialView, an open-source web browser-based interactive application for visualizing data and results from multiple 10x Genomics ST experiments. We anticipate SpatialView will be useful to a broad array of clinical and basic science investigators utilizing ST to study disease.

**Availability:** SpatialView is available at https://github.com/kendziorski-lab/SpatialView ; a demo application is available at https://www.biostat.wisc.edu/~kendzior/spatialviewdemo/

**Supplementary information:** Supplementary data are available at *Bioinformatics* online.

## 1 Introduction

A typical spatial transcriptomics (ST) experiment requires pre-processing, normalization, data integration, clustering, differential expression, other downstream analyses, and data visualization. A number of approaches exist for pre-processing and analyzing ST data including popular R packages such as Seurat (Hao Y *et al*., 2021), Giotto (Dries *et al*., 2021), and STUtility (Bergenstråhle *et al*.,2020), with similar applications such as Scanpy (Wolf *et al*., 2018) and Squidpy (Palla *et al*., 2022) available in Python. Graphical user interfaces (GUI) have also emerged including BBrowser (Le, T. *et al*., 2020), ST Viewer (Navarro *et al*., 2019), and SpatialDB (Zhen Fan *et al*., 2020). While these platforms have proven useful, there is still no convenient, intuitive tool for extensively and interactively visualizing data and results from ST experiments. The Loupe Browser from 10x Genomics as well as SpatialLIBD (Pardo *et al*., 2022) provide some capability toward this end. However, these platforms do not allow for an interactive analysis of multiple samples simultaneously, making it difficult to view and compare results across samples. The Loupe Browser also lacks general applicability as it relies on closed source code and is limited to results imported from Space Ranger. In addition, the available visualization tools lack salient analysis features including high resolution zoom in/out function for the images, in depth spot-by-spot browsing of marker genes and their function, and summary statistics for gene expression differences across groups. In short, there is currently no interactive visualization platform that allows users to easily study and compare results in an individual sample or across multiple ST samples. Toward this end, we developed SpatialView.

## 2 Methods

SpatialView enables fast, comprehensive, and interactive ST data visualization within and across multiple samples.

### 2.1 Input

SpatialView is a tool for visualizing results from 10x Genomics experiments and, as input requires the hematoxylin and eosin (H&E) strained images captured during ST sample preparation and the alignment files (scalefactors_json.json and tissue_positions_list.csv) from Space Ranger. The input files are stored in the application’s data directory. Results from downstream analyses such as clustering and identification of cell-population specific marker genes (or any genes of interest) are not required but can also be uploaded to enhance the visualization, but are not required. Unlike Loupe Browser, downstream analyses need not be carried out in Space Ranger because the complementary SpatialView packages SpatialViewR and SpatialViewPy (Supplementary file1, file2) can be used to import clustering results and genes of interest from R or Python-based pipelines, respectively. In addition, SpatialView does not technically require that clusters be determined by integrating data across samples; and unless comparative analyses across samples such as those described in Section 2.3 are not of interest, it is not necessary to integrate samples prior to clustering. However, if comparative analyses are of interest, SpatialView assumes that normalization and cluster identification have been done following sample integration.

Generally, the number of samples that can be handled in a single browser window is limited by the browser’s allocated memory capacity. Because SpatialView performs all the computations required for data visualization on the user’s browser, it does not require specialized server or memory management, eliminating back-and-forth server communications resulting in fast response times. In addition, since SpatialView requires only flat files for input, it may be used to visualize ST analysis results obtained from R, Python, Space Ranger, or any other data analysis platform, making it highly adaptable. Internally, SpatialView uses the D3, jQuery, Papa Parse, and Plotly opensource javascript libraries.

To visualize data from an experiment with 10 samples, for example, a computer with 4GB memory is suggested. While SpatialView can handle an expression matrix in csv format, we recommend the compressed sparse column-oriented (CSC) format to reduce data loading time and browser memory requirements; the sparse format is set as the default file export format in SpatialViewR and SpatialViewPy.

### 2.2 Visualization and analyses of individual samples

With the input files stored in the application’s data directory, SpatialView enables several interactive visualizations within a sample (Fig. 1) or across multiple samples (Fig. 2). The top panel of SpatialView shows H&E stained thumbnail images from all samples (Fig. 1A). For individual-sample visualization, a user can click to choose any sample from the top panel (Fig. 1B). The H&E stained tissue section with spot-specific cluster information is shown (Fig. 1C). For any spot selected on mouse-over (Fig. 1D), SpatialView provides a zoomed-in view to precisely locate that spot within the original tissue (Fig. 1E) along with spot-specific information such as cluster membership and expression of the top-five most highly expressed genes (Fig. 1F). A heatmap of cluster-specific marker gene expression (Fig. 1G) is also provided (if marker genes are specified in the input files) along with sample metadata such as tissue name and tissue type (Fig. 1H). Cluster-specific spots can be highlighted or hidden using the cluster buttons (Fig. 1I). A user can adjust spot-specific size and opacity, and H&E image brightness using the sliders (Fig. 1J). For easy access, marker genes for each cluster are listed at the right of the page (Figure 1K, 1L). A user can click on the gene name to view expression across the spots (Fig. S1); expression for any other gene can also be viewed by searching for that gene in the search box (Fig. 1M, Fig. S1). Detailed documentation from the *genecards*.*org* database can also be accessed by a single click for a gene in the heatmap or by a double click for a gene name in the list of markers.

**Figure 1:**
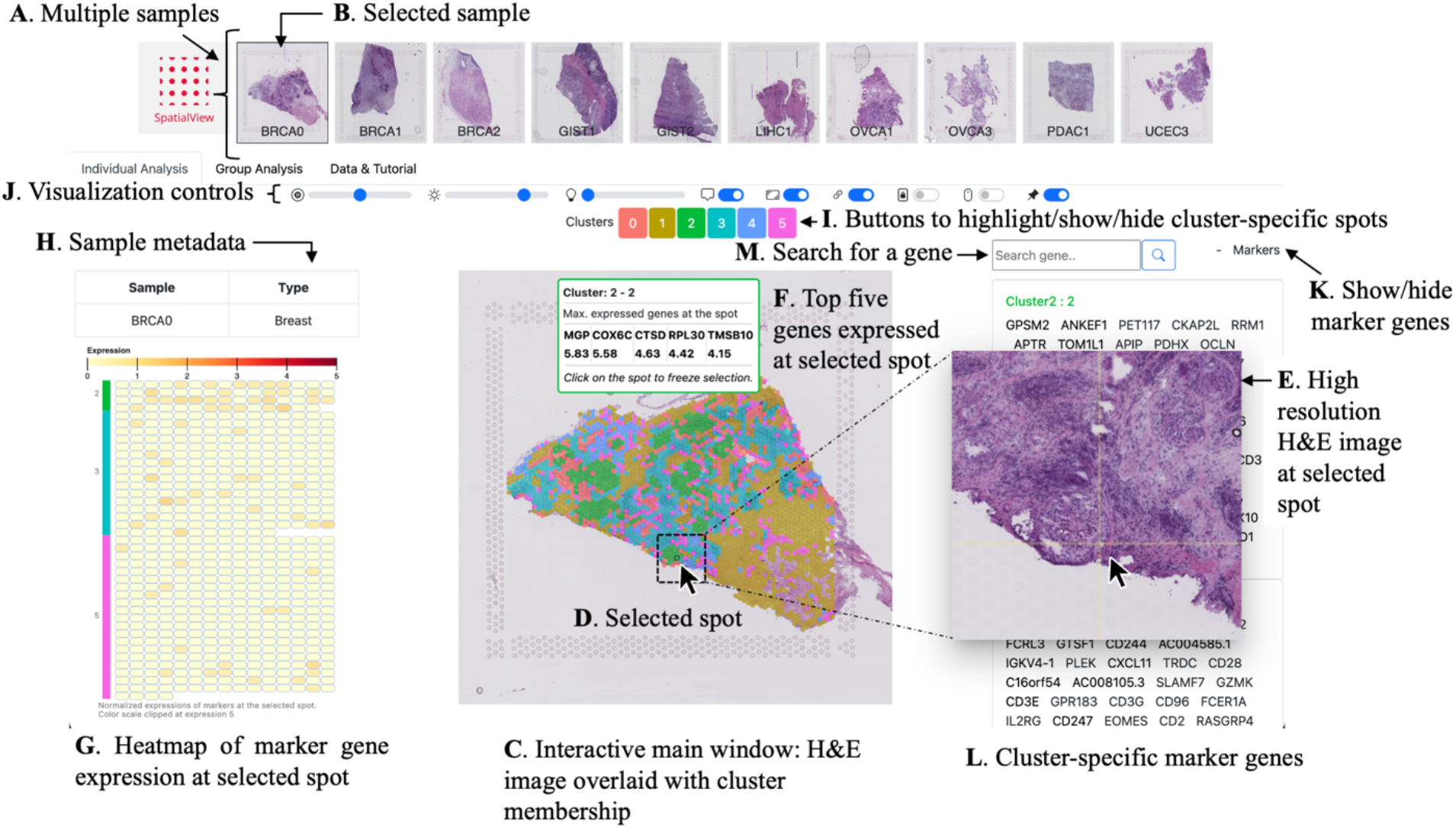
A screen shot of SpatialView. (A) The top panel of SpatialView shows H&E stain thumbnail images from all samples. (B) For single-sample visualization, a user may click to choose any sample from the top panel. For that sample, an H&E image with spot-specific cluster information is shown (C). Upon mouse over of any spot (D), SpatialView provides a zoomed-in H&E image (E) along with spot-specific information (cluster membership, top five gene expressions) (F) and a heatmap of marker gene expression at that spot (G). Sample metadata such as tissue name and tissue type are also shown (H). A user may choose to highlight, show or hide spots belonging to particular clusters by using the cluster-specific toggle buttons (I). Spot-specific size, opacity, and H&E image brightness can be adjusted using the sliders (J). If marker genes are provided, then they can be listed using the ‘markers’ link (K) and the cluster specific markers will be visible in the right panel (L). On click of a gene name or by searching a gene (M), the expression of the gene across the spots can be visualized (Fig. S1). Documentation about a gene in genecards.org can be accessed by a single click on a gene in the heatmap or by double click on a gene name in the list of markers.

**Figure 2:**
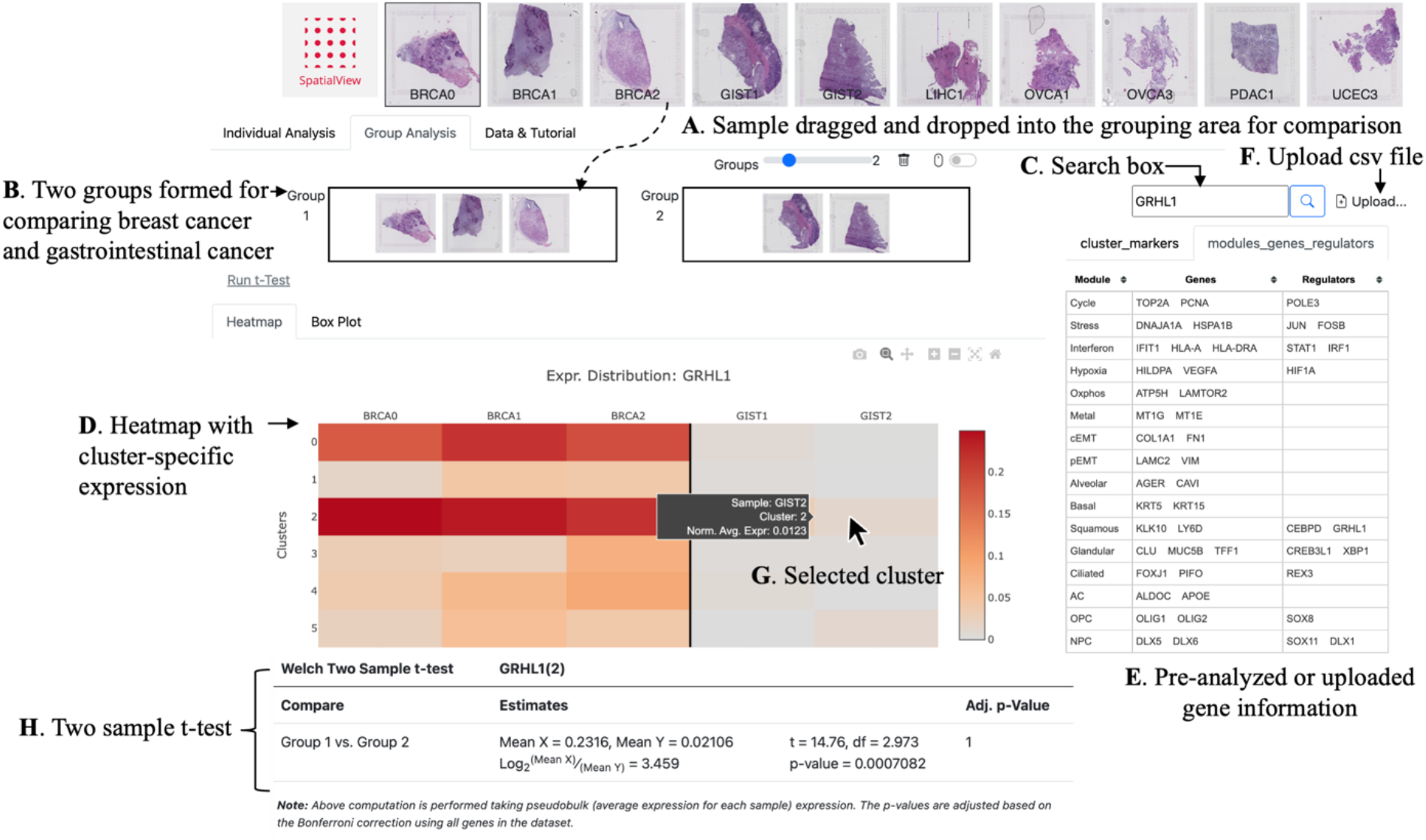
Visualization and analysis of multiple samples. The top panel of SpatialView shows H&E stain thumbnail images from all samples (A). In the ‘Group Analysis’ tab, samples can be dragged-and-dropped into groups (B). Here, two groups are shown (Supplementary Figure S2 shows visualization with four groups). A user may specify a gene of interest in the search box (C) and cluster-specific expression is displayed as a heatmap (D) or box plots (Figure S2). If a user has conducted an analysis to identify differentially expressed (DE) genes, or some other genes of interest, they can be presented by SpatialView in the right-side panel (E). For example, information from Fig. 1F (Barkley et al., 2022) is shown in the tab ‘modules_gene_regulators’. User can also upload (F) any csv file containing gene names and SpatialView will present it as a sortable table (Figure S2). On hovering over the heatmap rows (clusters), details about the sample specific average expression of the selected gene are shown (G), and a two-sample t-test is carried out (provided at least three samples are present in one group and two in another) (H).

Taken together, SpatialView allows a user to visually inspect cluster membership, expression of marker genes, gene function, and tissue morphology at any spot simultaneously.

### 2.3 Visualization and analyses of multiple samples

To compare gene expression patterns across multiple sample groups in an ST experiment (e.g., case vs. control, male vs. female, young vs. mid-age vs. old), a user of SpatialView can easily use the drag-and-drop feature in the group analysis tab. Figure 2 shows a simple two-group comparison with the groups created from the 10 total samples (Fig. 2A and 2B). SpatialView is not limited to two-group comparisons. Section 3 and Supplementary Figure S2 show results from a four-group analysis; Figure S3 shows results from a one-group analysis where comparisons across clusters instead of across samples are of interest. After forming the groups and searching for a gene of interest in the search box (Fig. 2C), a user can view a heatmap and boxplots of cluster-specific average expression across the samples for any specified gene (Fig. 2D, Fig. S2G). As noted in Section 2.1, SpatialView assumes that the samples have been normalized and integrated prior to identification of clusters to ensure comparable expression across samples.

If a user has conducted an analysis to identify differentially expressed (DE) genes, or has a list of other genes of interest, a csv file with those lists can be uploaded (Fig. 2F) and is presented in SpatialView as a sortable table (Fig. 2E, S2). Comparative visualizations are generated by clicking on any gene in the uploaded list (Fig. S2E). Cluster-specific two-sample t-tests can also be conducted for a selected gene by hovering over any row (cluster) in the heatmap (Fig. 2G); results are shown below the heatmap (Fig. 2H).

### 2.4 Testing

With two groups of samples selected as in Fig. 2B, SpatialView can be used to conduct cluster-specific pseudobulk t-tests for a specified gene by hovering over the gene’s heatmap (Fig. 2G). The cluster-specific pseudobulk t-test is a t-test conducted on expression averaged across all cluster-specific spots in a sample; tests require at least three samples in one group and two in another. When there are multiple groups, all pairwise comparisons are performed. Genome-wide testing can be performed using the ‘Run t-Test’ link (Fig. S2B). To limit the test results, additional filters can be used (Fig S2C). Note that genome-wide testing on a browser may take a few minutes. Reported p-values are adjusted based on the Bonferroni correction using the total number of genes present in the dataset. When there is only one group of samples defined, a pseduobulk t-test can be performed between a selected cluster (selected on mouse-over) vs. the rest of the clusters (Fig. S3).

### 2.5 Metadata, Data Sharing, and Tutorial

The metadata for each sample can be accessed along with single sample visualizations (Fig. 1H). Additional information about the samples or about the experiment can be annotated in the ‘Data & Tutorial’ tab (above Visualization controls, Fig. 1J). An investigator can host SpatialView to allow visualization of their data by anyone with access to a web browser. Details are provided for this and other features of SpatialView in a video tutorial accessible from the application.

## 3 Use Case

To illustrate how SpatialView can be used to characterize clusters, to investigate marker gene expression, and to perform comparative analysis among group of samples, we considered ten ST samples from a study of cancer (3 breast, 2 gastrointestinal, 1 liver, 2 ovarian, 1 pancreas, and 1 endometrium) detailed in Barkley *et al*., 2022. As described in Section 2.1, SpatialView expects H&E strained tissue images and alignment files for each sample. Since we will illustrate how SpatialView can be used to investigate spatial patterns of marker genes within and across clusters, we also uploaded results from clustering and DE testing.

After following the Seurat data analysis pipeline in R for pre-processing, normalization, integration, and clustering, six clusters were identified. For each cluster, one vs. rest pseudobulk t-tests were performed and DE genes were defined as marker genes for each cluster (methods for pre-processing, cluster identification, and DE testing are specified in supplementary file1). Finally, the processed data were exported to SpatialView using the *prepare10x_from_seurat* function in the SpatialViewR package.

Fig. 1A shows the H&E stained tissue images of each sample at the top of SpatialView’s application page. A breast cancer sample BRCA0 is selected (Fig. 1B) and loaded into the main window (Fig. 1C). The spots of the sample are laid over the histology image and are color-coded to match the corresponding cluster button colors (Fig. 1I). Suppose we are interested in identifying clusters mostly comprised of tumor spots as well as those mostly comprised of normal spots. To do this, we can zoom into the H&E image by hovering on any spot (Fig. 1D); the zoomed-in H&E image will be shown with the mouse pointer at the chosen spot (Fig. 1E). Repeating this a few times, we can visually confirm that cluster 2 is almost entirely made up of tumor spots. Spots belonging to cluster 2 can be isolated using the buttons that highlight/hide cluster-specific spots (Fig. 1I and Fig. S1C). Our prior analysis identified the PRELI domain containing 3B gene *PRELID3B* as a marker gene for cluster 2, and so it is of interest to visualize expression of this gene across the spots of cluster 2. After searching for the *PRELID3B* gene in the search box (Fig. S1D), expression across the selected spots can be visualized in the interactive main window. Fig. S1E shows *PRELID3B* expression across spots in clusters 2 and 3; cluster 2 (tumor) shows slightly higher expression of this marker gene compared to cluster 3.

For comparative analysis among the four cancer-types (breast, gastrointestinal, liver, and ovarian), four groups are made by dragging-and-dropping the corresponding samples as shown in Fig. S2A. A genome-wide psuedobulk t-test is conducted (Fig. S2B) for genes having at least 5 samples, with at least 3 in one group (Fig. S2C); a results file is generated and uploaded into the application by the user (Fig. S2D). After uploading the results file, an additional tab with a sortable table is added in the right-side panel (Fig. S2D). SpatialView automatically detects the gene names from the uploaded file and allows them to be searched interactivity. In the four-group analysis, for example, we observe in cluster 2 that *BMPR1B*, a member of the threonine kinase receptor family, has higher expression in group one as compared to group two (Fig. S2D). For visual inspection, the gene is selected (Fig. S2E) and a heatmap showing average expression in all the selected samples is loaded with the samples in the heatmap arranged to match the groupings. Fig. S2F shows higher expression of *BMPR1B* in the breast cancer samples; the box plots confirm that the observation is not due to outliers (Fig. 2SG). This finding suggests a role for *BMPR1B* in discriminating cancer types that warrants further investigation.

## 4 Summary

SpatialView is an interactive web application designed to visualize ST data and results including across multiple samples. SpatialView can be used to visualize results directly from Space Ranger, or can be easily accessed from the R and Python environments using the SpatialViewR and SpatialViewPy packages. The application can be run on a local computer or on a remote web server. SpatialView’s interactive design, coupled with drag-and-drop features, allows users to effectively visualize gene expression patterns across multiple ST samples and cell types.

## Supporting information

https://kendziorski-lab.github.io/projects/spatialview/SpatialView_Tutorial_Using_Seurat.html

https://github.com/kendziorski-lab/SpatialViewPy/blob/main/notebooks/tutorial.ipynb

---

**Figure S1:**
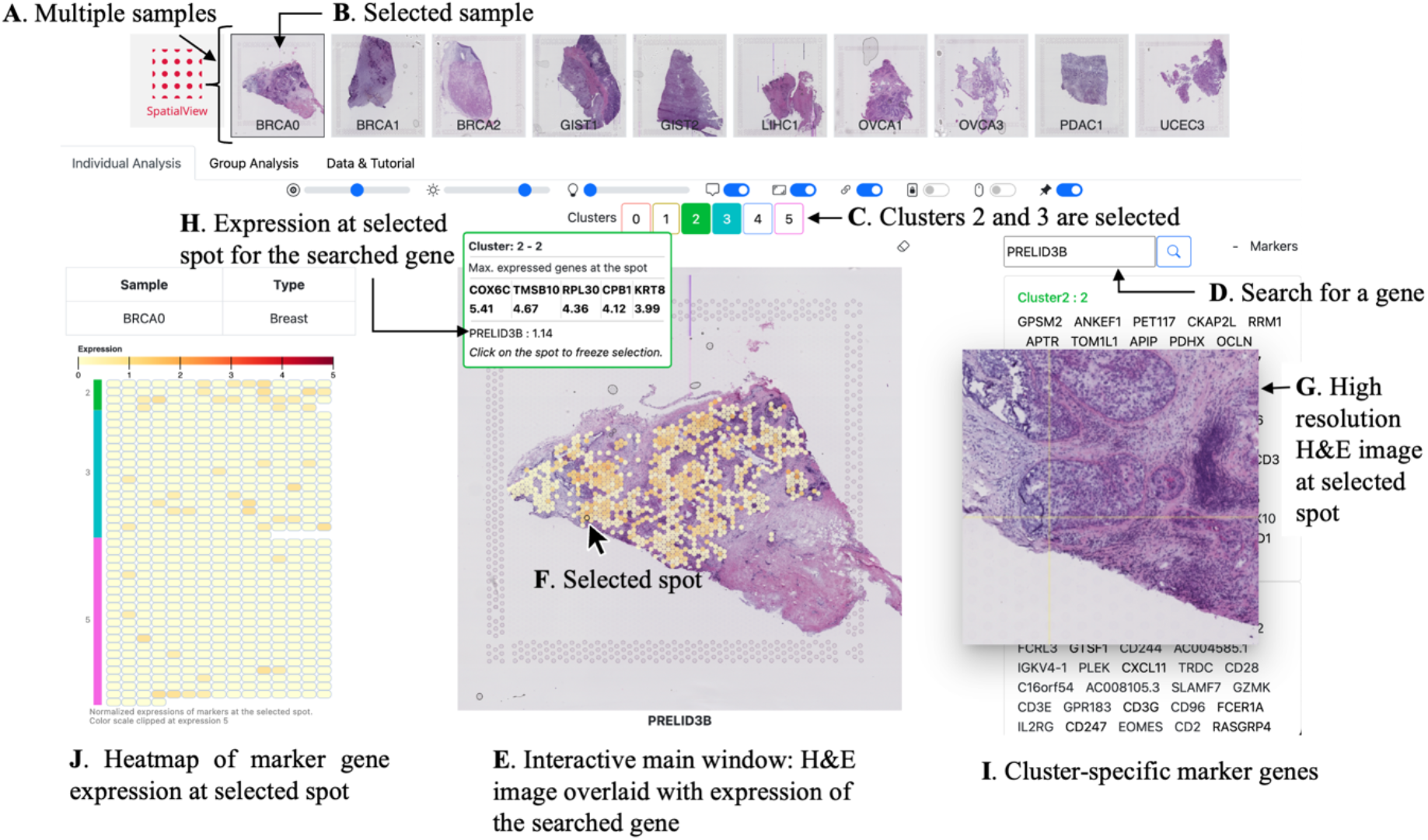
A screen shot of SpatialView. The top panel of SpatialView shows H&E stained thumbnail images from all samples (A). For single-sample visualization, a user may click to choose any sample from the top panel (B). A user can choose interested cluster specific spots using the toggle buttons (C). After searching a gene name in the search box (D), its expression across the selected cluster specific spots will be shown in the interactive panel (E). On mouse over a spot (F), a zoomed-in H&E image is shown (G); spot-specific information includes expression of the searched gene at the selected spot (H). Marker genes are shown in the right-side panel (I) and heatmap showing expressions of markers at the selected spot is shown in the left-side panel (J).

**Figure S2:**
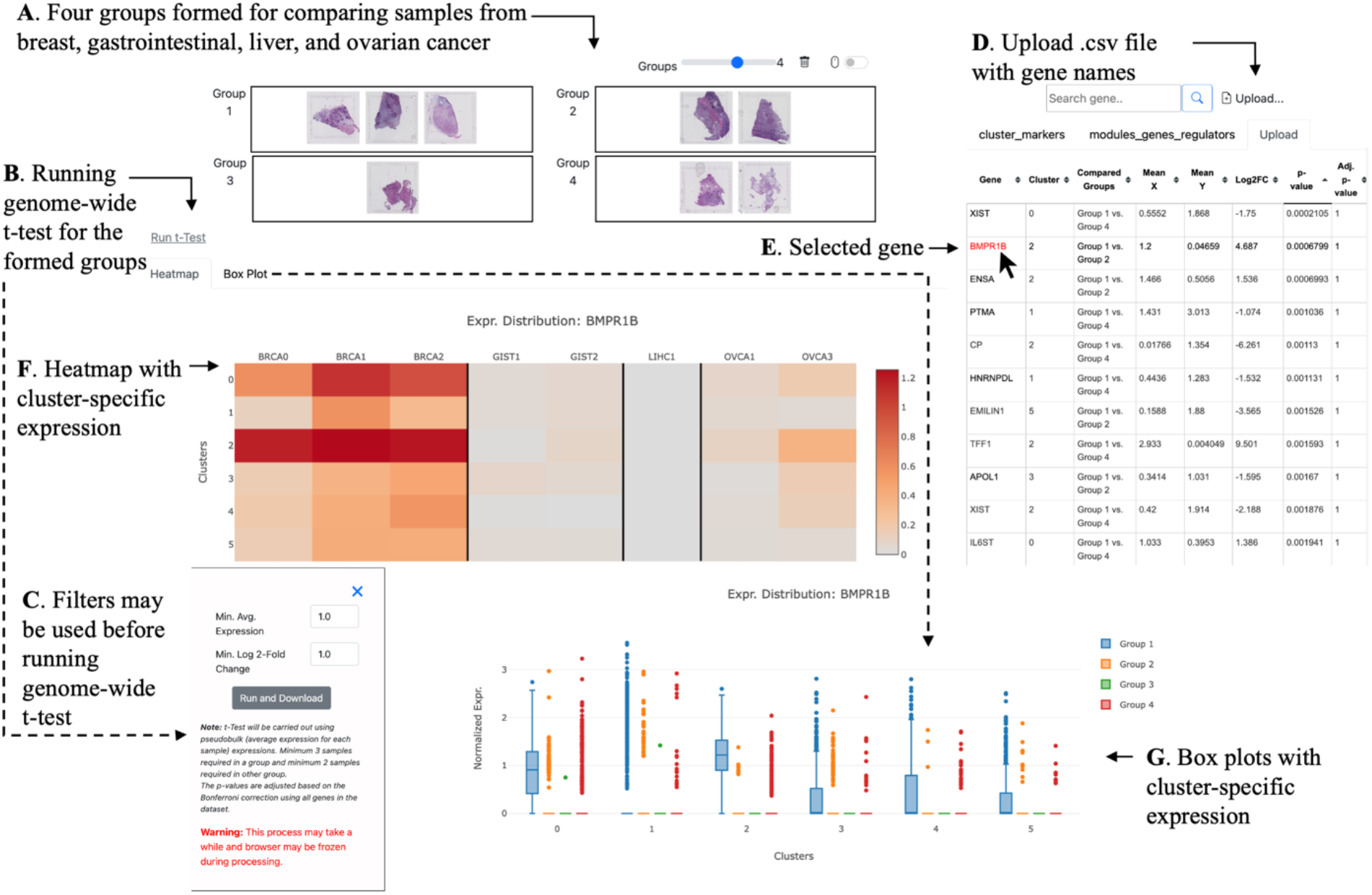
Four-group comparative analysis. Four groups are formed representing four cancer types (breast, gastrointestinal, liver, and ovarian cancer) by dragging and dropping the corresponding samples into the grouping area (A). After forming the groups, a user may search for a gene of interest in the search box to visualize that gene’s expression in any cluster, as described in Fig. 2. Additionally, instead of investigating individual genes, a user may initiate a genome-wide pairwise t-test using the ‘Run t-Test’ link (B). Before running the test, additional filters may be applied to refine the set of genes considered (C). After completion of the tests, a results file will be saved; for visualizing results, the file can be uploaded using the upload link (D). After uploading the results file, an additional tab with a sortable table will appear in the right-side panel. SpatialView automatically detects the gene names from the uploaded file and allows them to be searched interactivity. On mouse over of the *BMPR1B* (E), we see in the table that its expression is higher in group 1 than in group 2 for cluster 2 (log 2-fold change 4.687); cluster wise expressions of *BMPR1B* across the groups is also shown in a heatmap (F) and box plots (G).

**Figure S3:**
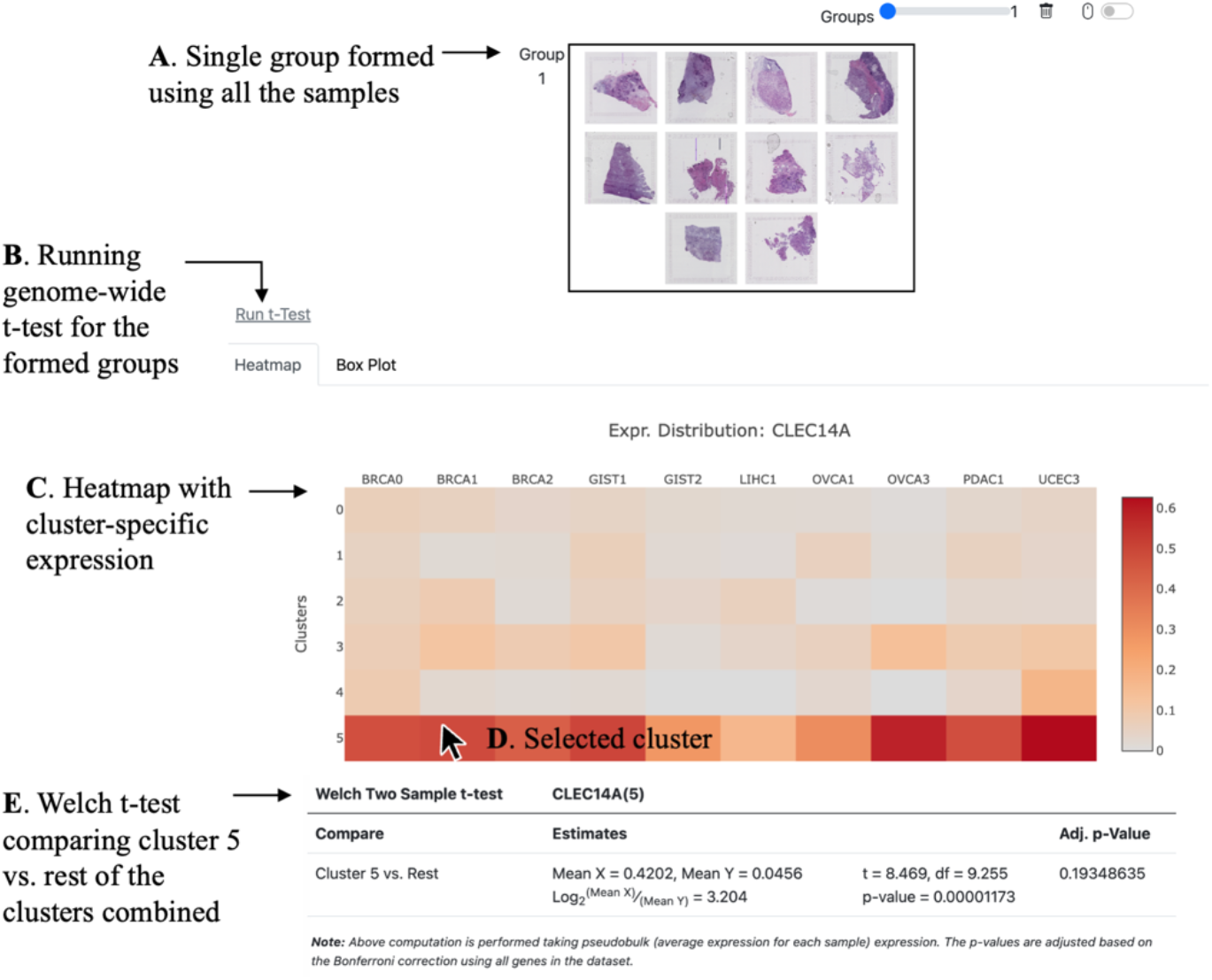
One-group analysis with comparisons across clusters. For this analysis, a single group is formed and samples of interest are dragged-and-dropped into the grouping the area (A). Once the group is formed, a user may search for a gene of interest (not shown here) or perform a genome-wide search as described in Figure 2 and Figure S2. In the case of one-group, when genome-wide t-tests are carried out using the ‘Run t-test’ link (B), tests are conducted for one cluster vs. the rest of the clusters. This is also the case on mouse-over of any row (cluster) of the heatmap (C, D).

