## Supplementary material for "SpatialView: An interactive web application for visualization of multiple samples in spatial transcriptomics experiments": https://kendziorski-lab.github.io/projects/spatialview/SpatialView_Tutorial_Using_Seurat.html: Supplementary_file1_seurat_tutorial.html

SpatialView Tutorial: Exporting data from Seurat


### SpatialView Tutorial: Exporting data from Seurat

###### Chitrasen Mohanty

#### 06/01/2023

In this tutorial we are using Spatial Transcriptomics (ST) data
published in (Barkley et al. 2022). This
data contains multiple samples from different cancer types, such as
breast, gastrointestinal, liver, ovary, pancreas, endometrium, and
others. This data is helpful to understand the heterogeneity in tumor
micro environment (TME) among different cancer types. For this
demonstration, we will be using the ten samples (3 breast, 2
gastrointestinal, 1 liver, 2 ovarian, 1 pancreas, and 1 endometrium)
generated using 10x SpaceRanger. We will use Seurat (Hao et al. 2021) for data pre-processing and
integrating the samples. With the integrated data, we will do clustering
followed by differential expression (DE) analysis to identify the marker
genes of the clusters. Finally, we will export the analyzed data to
**SpatialView** (Mohanty *et al*., 2023) for
interactive visualization.

*Note that it requires to have python installed in your system.
Python comes with many operating system pre-installed (such as Linux,
Mac OS). To install python, please follow https://www.python.org/doc/.*

Tutorial for SpatialView
using SpatialExperiment

```
#Install SpatialviewR if not yet installed.
remotes::install_github("kendziorski-lab/SpatialviewR")

library(stringr)
library(dplyr)
library(Seurat)
library(R.utils)
library(ggplot2)
library(SpatialViewR)

# Installing a few additional packages for matrix operations for DE analysis
install.packages("grr")
install.packages("https://cran.r-project.org/src/contrib/Archive/Matrix.utils/Matrix.utils_0.9.8.tar.gz", type = "source", repos = NULL)

library("Matrix.utils")
```

### Data

The tar file containing all the required samples can be downloaded
from GEO (https://www.ncbi.nlm.nih.gov/geo/query/acc.cgi?acc=GSE203612).
Note that, the file size is 675MB. Once the data is downloaded to the
local computer, the following code untars the contents and places them
in their appropriate sub directories.

```
#provide the correct path to the downloaded tar file
data_path = "../../data_others/spatial/cancer_multi/GSE203612_RAW.tar"
set.seed(2023)
dir_name = dirname(data_path)
base_name = str_split(basename(data_path), pattern = "\\.")[[1]][1]
untar(data_path, exdir = file.path(dir_name, base_name))
data_path = file.path(dir_name, base_name)

#If the files are already uncompressed then change the path in the below line.
file_names <- list.files(data_path)
files.df <- as.data.frame(str_split(file_names, "_", n = 4, simplify = TRUE))
colnames(files.df) <- paste0("col",1:4)
files.df <- files.df %>% rowwise() %>% mutate(sample_name = ifelse(col2 == "NYU", col3, col2))
files.df$file_name = file_names

#currently we are using NYU samples only.
for (i in 1:nrow(files.df)) {
  target_file_name <- files.df[i, "file_name"]
  if (str_detect(target_file_name, "Vis") & files.df[i, "col2"] == "NYU") {
    
    target_file_name <- str_remove(target_file_name, pattern = ".*processed_")
    output_dir <- file.path(data_path, files.df[i,"sample_name"])
    if (str_detect(target_file_name, "spatial")) {
      output_dir <- file.path(output_dir,"spatial")
      target_file_name <- str_remove(target_file_name, pattern = ".*spatial_")
    }
    
    if (!dir.exists(output_dir)) {dir.create(output_dir, recursive = TRUE)}
    
    file.copy(from = file.path(data_path, files.df[i, "file_name"]), 
              to = file.path(output_dir, target_file_name), 
              overwrite = TRUE)
    #if the file is compressed then decompress it
    if (str_ends(target_file_name, ".gz")) {
      gunzip(file.path(output_dir, target_file_name), overwrite = TRUE)
    }
  }
   
    if(!isDirectory(file.path(data_path, files.df[i, "file_name"]))){
    file.remove(file.path(data_path, files.df[i, "file_name"]))
  }
}
```

#### Reading data as Seurat object:

```
#confirm the data_path
data_path = "../../data_others/spatial/cancer_multi/GSE203612_RAW"

TME_10x.list <- lapply(list.files(data_path), function(d){
  data.sample <- Load10X_Spatial(data.dir = file.path(data_path, d), 
                                slice = d)
  data.sample$orig.ident <- d
  data.sample <- NormalizeData(data.sample, assay = "Spatial", verbose = FALSE)
  data.sample
})
names(TME_10x.list) <- list.files(data_path)
```

### Data Processing

#### Pre-processing

As a quality control step, spots with fewer than 500 UMIs or more
than 30% mitochondrial or ribosomal reads were filtered out.

We filter out genes having present in less than 2 samples or less
than 15 spots in the available samples.

```
data_cleaning <- function(data.list,min_depth=500, mt.pct.max = 30, rib.pct.mx = 30, min_cell_for_gene = 15,
                          min.samples = 2){
  data.list <- lapply(data.list, function(x){
    x$percent.mt <- PercentageFeatureSet(x, pattern = "^MT-")
    x$percent.rp <- PercentageFeatureSet(x, pattern = "^RP[SL]")
    subset(x, nCount_Spatial >= min_depth & percent.mt <= mt.pct.max & percent.rp <= rib.pct.mx)
  })
  
  g_in_cells_counts = matrix(nrow=nrow(data.list[[1]]), ncol = length(data.list))
  for(i in seq_along(data.list)){
    #print(dim(data.list[[i]]))
    g_in_cells_counts[,i] <- Matrix::rowSums(data.list[[i]]@assays$Spatial@counts > 0) >= min_cell_for_gene
  }
  
  rownames(g_in_cells_counts) <- rownames(data.list[[1]])
  g_in_cells_counts <- g_in_cells_counts[rowSums(g_in_cells_counts) >= min.samples,]
  sel_genes <- rownames(g_in_cells_counts)
  
  data.list <- lapply(data.list, function(x){
    subset(x, features = sel_genes)
  })
  
  return(data.list)
}

TME_10x.list <- data_cleaning(TME_10x.list)
```

#### Normalization

We are performing logNomalization using *NormalizeData*
function from Seurat.

```
TME_10x.list.norm <- lapply(X = TME_10x.list, FUN = function(x) {
    NormalizeData(x, assay = "Spatial", verbose = FALSE)
})
```

#### Integration Using Seurat Pipeline

Now, we will integrate all the samples using Seurat data integration
pipeline.

```
#getting 10% of the genes as HVG
features <- SelectIntegrationFeatures(object.list = TME_10x.list.norm, 
                                      nfeatures = 1650, verbose = FALSE)

anchors <- FindIntegrationAnchors(object.list = TME_10x.list.norm,
                                  normalization.method = "LogNormalize",
                                  anchor.features = features, dims = 1:30,
                                  l2.norm = TRUE,k.anchor = 5, k.filter = 200,
                                  max.features = 200, n.trees = 50,
                                  verbose = FALSE, 
                                  )
integrated.data  <- IntegrateData(anchorset = anchors,
                                   new.assay.name = "integrated",
                                   normalization.method = "LogNormalize", 
                               k.weight = 100, verbose = FALSE)
```

#### Clustering

With the integrated data, we will perform clustering. Note that, the
number of clusters detected is sensitive to the input parameters used in
the functions. The parameters used in this example are for demonstration
purpose.

```
DefaultAssay(integrated.data) <- "integrated"
integrated.data <- ScaleData(integrated.data, verbose = FALSE)
integrated.data <- RunPCA(integrated.data, verbose = FALSE)
integrated.data <- RunUMAP(integrated.data, assay = "integrated", dims = 1:30, verbose = FALSE,
                           min.dist = 0.01, spread = 3) %>%
              FindNeighbors(verbose = FALSE) %>%
              FindClusters(resolution = 0.2, n.start = 25, n.iter = 25, 
                           algorithm = 1,
                           verbose = FALSE)
```

After this step, we will be observing six clusters (Numbered from 0
to 5).

```
library(ggplot2)
g <- DimPlot(integrated.data, reduction = "umap", label = TRUE, repel = TRUE, 
        shuffle = TRUE)+ggtitle("UMAP by Cluster")
g <- g + theme(axis.line=element_blank(),axis.text.x=element_blank(),
          axis.text.y=element_blank(),axis.ticks=element_blank(),
          axis.title.x=element_blank(),
          axis.title.y=element_blank())

g
```

#### Differential Expression (DE) Analysis

Finding marker genes in each cluster we will perform t-test comparing
each cluster with rest of the clusters using pseudobulk (average
expression for each sample) expressions.

*This step is optional, you may use other DE analysis as
well.*

```
DefaultAssay(integrated.data) <- "Spatial"

# DE analysis
all.markers.pseudoBulk.tTest <- function(object = integrated.data, 
                              assay = "Spatial", slot = "data",
                              only.pos = FALSE, log2FC.min = 0){
  
  groups <- object@meta.data[, c("seurat_clusters", "orig.ident")]
  aggr_sum <- aggregate.Matrix(t(GetAssayData(object = object,
                                                 assay = "Spatial",
                                                 slot = "data")), 
                                  groupings = groups, fun = "sum")
  
  aggr_num <- aggregate.Matrix(t(GetAssayData(object = object,
                                                 assay = "Spatial",
                                                 slot = "data") > -1), 
                                  groupings = groups, fun = "sum")

  aggr_counts = aggr_sum/aggr_num

  aggr_counts <- t(aggr_counts)
  cluster_names <- unique(object@meta.data$seurat_clusters)
  test_res.df_all <- lapply(cluster_names, function(c){
    matched_cols <- str_detect(base::colnames(aggr_counts), paste0("^", c,"_"))
    
    aggr_counts.sub <- aggr_counts
    
    fc <- log2(rowMeans(aggr_counts.sub[,matched_cols])) -  log2(rowMeans(aggr_counts.sub[,!matched_cols]))
    if(only.pos){
      aggr_counts.sub <- aggr_counts.sub[fc > 0,]
      fc <- log2(rowMeans(aggr_counts.sub[,matched_cols])) -  log2(rowMeans(aggr_counts.sub[,!matched_cols]))
    }
    if(log2FC.min > 0){
      aggr_counts.sub  <- aggr_counts.sub[abs(fc) > log2FC.min,]
    }
    
    test_res <- lapply(rownames(aggr_counts.sub), function(g){
      res <-  t.test(aggr_counts.sub[g,matched_cols], aggr_counts.sub[g,!matched_cols])
      return(c(res$estimate[1], res$estimate[2], res$p.value))
    })
    test_res.df <- data.frame(matrix(unlist(test_res), 
                                     nrow = nrow(aggr_counts.sub), 
                                     byrow = T))
    base::colnames(test_res.df) <- c("Mean.X", "Mean.Y", "p_value")
    test_res.df$Log2FC <- log2(test_res.df[,"Mean.X"]) - log2(test_res.df[,"Mean.Y"])
    test_res.df$Cluster = c
    test_res.df <- test_res.df[,c("Cluster","Mean.X", "Mean.Y", "Log2FC", "p_value")]
    test_res.df$p_val_adj <- p.adjust(test_res.df$p_value, method = "BH")
    test_res.df$Gene <- rownames(aggr_counts.sub)
    rownames(test_res.df) <- rownames(aggr_counts.sub)
    test_res.df
  }) 
  return(do.call("rbind", test_res.df_all))
}

markers.bulkDE <- all.markers.pseudoBulk.tTest(object = integrated.data, 
                              assay = "Spatial", slot = "data", only.pos = TRUE,
                              log2FC.min = 1)

markers.bulkDE3 <- all.markers.pseudoBulk.tTest(object = integrated.data,
                              assay = "Spatial", slot = "data", only.pos = TRUE,
                              log2FC.min = 1)


selected.markers.bulkDE <- markers.bulkDE %>% 
  filter(p_val_adj < 0.1) %>% 
  arrange(desc(Log2FC))

selected.markers.list.bulkDE <- lapply(levels(integrated.data@meta.data$seurat_clusters), function(c){
  selected.markers.bulkDE$Gene[selected.markers.bulkDE$Cluster == c]
})
```

### Visualization Using SpatialView

```
#confirmt the export path
export_path = "outs/TME_SpatialView/"
dir.create(export_path, recursive = TRUE)

#adding sample information to the visualization
sampleInfo <- data.frame(sample = c("BRCA0", "BRCA1", "BRCA2", 
                                    "GIST1", "GIST2", 
                                    "LIHC1", "OVCA1",  "OVCA3",  
                                   "PDAC1",   "UCEC3"),
                         type = c("breast", "breast", "breast",
                                  "gastrointestinal", "gastrointestinal",
                                  "liver", "ovary", "ovary",
                                "pancreas", "endometrium"))

data.dir <- file.path(data_path, list.files(data_path))

prepare10x_from_seurat(integrated.data, data.dir, 
                       export.path = export_path, 
                       clusterCol = "seurat_clusters",
                       projectName = "TME",
                       downloadRepo = TRUE,
                       exprRound = 2,
                       clusterGenes = selected.markers.list.bulkDE,
                       sampleInfo = sampleInfo,
                       verbose = TRUE)
```

The SpatialView application will be launched in your preferred web
browser.

If you are interested to evaluate one or multiple sets of genes in
the group analysis, you can easily do so by placing the files in the
group\_data/group\_genes directory. This file should be a csv file with
following columns “cluster”,“color”,“name”,“genes”. The “genes” column
contains all the genes with ‘,’ separated.

For example

```
copyFile("outs/TME_SpatialView/TME/data/BRCA0/cluster_info.csv", "outs/TME_SpatialView/TME/group_data/group_genes/")
```

You can also save multiple csv files containing genes (no additional
formatting required) in the group\_data/show\_tables directory.
SpatialView automatically shows this data as an interactive table.

After adding the files to the group\_genes directory, use
reload/refresh button on the web page to reload the application.
