## Supplementary material for "SpatialView: An interactive web application for visualization of multiple samples in spatial transcriptomics experiments": https://github.com/kendziorski-lab/SpatialViewPy/blob/main/notebooks/tutorial.ipynb: Supplementary_file2_scanpy_tutorial.html


In [1]:

```
!pip install spatialviewpy
```

May need to install following additional packages.

In [2]:

```
## If running in conda environment
# !conda install -c conda-forge python-annoy -y

# !pip install scanorama
# !pip install requests
```

In [3]:

```
import scanpy as sc
import anndata as an
import pandas as pd
import numpy as np
import matplotlib as mpl
import matplotlib.pyplot as plt
import seaborn as sns
import scanorama

from spatialviewpy import prepare10x_from_scanpy

from os import listdir, path
from collections import OrderedDict

import warnings
warnings.filterwarnings('ignore')
```

In [4]:

```
# import numpy as np
# from scipy.io import mmwrite
# from zipfile import ZipFile
# import seaborn as sns
# import requests
# import os
# import shutil
# import json
# import gzip
```

In this tutorial we are using Spatial Transcriptomics (ST) data published in (Barkley et al. 2022). This data contains multiple samples from different cancer types, such as breast, gastrointestinal, liver, ovary, pancreas, endometrium, and others. This data is helpful to understand the heterogeneity in tumor micro environment (TME) among different cancer types. For this demonstration, we will be using the ten samples (three breast, two gastrointestinal, one liver, two ovarian, one pancreas, and one endometrium) generated using 10x SpaceRanger. We will use Scanpy (Wolf *et al*., 2018) for data pre-processing and Scanorama (Hie *et al*., 2019) for integrating the samples. With the integrated data, we will do clustering followed by differential expression (DE) analysis to identify the marker genes of the clusters. Finally, we will export the analyzed data to SpatialView (Mohanty et al., 2023) for interactive visualization. Note that analysis steps are for demonstration purpose only.

#### Loading data¶

In [5]:

```
#path to data directory
data_path = "../../data_others/spatial/cancer_multi/GSE203612_RAW/"

sample_names = listdir(data_path)
sample_names = [d for d in sample_names if d[0] != '.']
sample_names = np.sort(sample_names)

sample_names
```

Out[5]:

```
array(['BRCA0', 'BRCA1', 'BRCA2', 'GIST1', 'GIST2', 'LIHC1', 'OVCA1',
       'OVCA3', 'PDAC1', 'UCEC3'], dtype='<U5')
```

In [6]:

```
samples_dict = OrderedDict()
for samp in sample_names:
    adata = sc.read_10x_h5(path.join(data_path, samp, 'filtered_feature_bc_matrix.h5'))
    adata.var_names_make_unique()
    adata.var["mt"] = adata.var_names.str.startswith("MT-")
    adata.var["rp"] = adata.var_names.str.startswith("RP[SL]")
    sc.pp.calculate_qc_metrics(adata, qc_vars=["mt","rp"], inplace=True)
    samples_dict[samp] = adata
```

```
OMP: Info #271: omp_set_nested routine deprecated, please use omp_set_max_active_levels instead.
```

In [7]:

```
sample_info = pd.DataFrame({'sample': ["BRCA0", "BRCA1", "BRCA2", "GIST1",  
                                   "GIST2", "LIHC1", "OVCA1",  "OVCA3",  
                                   "PDAC1",   "UCEC3"],
                         'tissue' : ["breast", "breast", "breast", "gastrointestinal",
                                "gastrointestinal", "liver", "ovary", "ovary",
                                "pancreas", "endometrium"]}
                         )
sample_info
```

Out[7]:

|  | sample | tissue |
| --- | --- | --- |
| 0 | BRCA0 | breast |
| 1 | BRCA1 | breast |
| 2 | BRCA2 | breast |
| 3 | GIST1 | gastrointestinal |
| 4 | GIST2 | gastrointestinal |
| 5 | LIHC1 | liver |
| 6 | OVCA1 | ovary |
| 7 | OVCA3 | ovary |
| 8 | PDAC1 | pancreas |
| 9 | UCEC3 | endometrium |

#### Preprocessing and Normalization:¶

As a quality control step, spots with fewer than 500 UMIs or more than 30% mitochondrial or ribosomal reads were filtered out. Also, we are removing genes which has less than 5 num of spots.

Followed by data preprocessing, data is normalized.

In [8]:

```
for name, adata in samples_dict.items():
    sc.pp.filter_cells(adata, min_counts=500)
    adata = adata[adata.obs["pct_counts_mt"] < 30]
    adata = adata[adata.obs["pct_counts_rp"] < 30]
    sc.pp.filter_genes(adata, min_cells=5)
    
    sc.pp.normalize_total(adata, target_sum = 10000, inplace=True, copy= True)
    sc.pp.log1p(adata)
    sc.pp.highly_variable_genes(adata, flavor="seurat", n_top_genes=2000, inplace=True)
    samples_dict[name] = adata
```

In [9]:

```
# We will use the unscaled normalized data for data visualization, thus saving them in a layer
adata_norm = sc.concat(
    list(samples_dict.values()),
    label="sample_id",
    uns_merge="unique",
    index_unique="_",
    keys = samples_dict.keys()
)
```

#### Data Integration¶

Data integrated using Scanorama.

In [10]:

```
adatas_cor = scanorama.correct_scanpy(list(samples_dict.values()), 
                                      return_dimred=True,
                                     sigma = 10)
```

```
Found 11047 genes among all datasets
[[0.         0.26403642 0.39789196 0.03920724 0.00131752 0.01668863
  0.16022727 0.00249169 0.02736318 0.03952569]
 [0.         0.         0.01820941 0.52200303 0.0030349  0.00856269
  0.01365706 0.05766313 0.12746586 0.03793627]
 [0.         0.         0.         0.0353296  0.         0.00214684
  0.08181818 0.00249169 0.05659204 0.00382848]
 [0.         0.         0.         0.         0.35341365 0.01406728
  0.00511364 0.08388704 0.18843284 0.02197329]
 [0.         0.         0.         0.         0.         0.
  0.         0.         0.00808458 0.00229709]
 [0.         0.         0.         0.         0.         0.
  0.00568182 0.09302326 0.         0.00229709]
 [0.         0.         0.         0.         0.         0.
  0.         0.00083056 0.10447761 0.19772727]
 [0.         0.         0.         0.         0.         0.
  0.         0.         0.00166113 0.78571429]
 [0.         0.         0.         0.         0.         0.
  0.         0.         0.         0.06032338]
 [0.         0.         0.         0.         0.         0.
  0.         0.         0.         0.        ]]
Processing datasets (7, 9)
Processing datasets (1, 3)
Processing datasets (0, 2)
Processing datasets (3, 4)
Processing datasets (0, 1)
Processing datasets (6, 9)
Processing datasets (3, 8)
Processing datasets (0, 6)
Processing datasets (1, 8)
Processing datasets (6, 8)
```

In [11]:

```
samples_dict.keys()
```

Out[11]:

```
odict_keys(['BRCA0', 'BRCA1', 'BRCA2', 'GIST1', 'GIST2', 'LIHC1', 'OVCA1', 'OVCA3', 'PDAC1', 'UCEC3'])
```

We will concatenating the two dataset with uns\_merge="unique" strategy, in order to keep both images from the visium datasets in the concatenated anndata object following the step https://scanpy-tutorials.readthedocs.io/en/latest/spatial/integration-scanorama.html

In [12]:

```
adata_spatial = sc.concat(
    adatas_cor,
    label="sample_id",
    uns_merge="unique",
    index_unique="_",
    keys = samples_dict.keys()
)
```

In [13]:

```
sc.pp.neighbors(adata_spatial, use_rep="X_scanorama")
sc.tl.umap(adata_spatial)
sc.tl.leiden(adata_spatial,resolution=0.15, key_added="clusters")

sc.pl.umap(
    adata_spatial, color=["clusters", "sample_id"], palette=sc.pl.palettes.default_20
)
```

In [14]:

```
adata_spatial.layers['normalized_counts'] = adata_norm.X
```

#### DE test¶

In [15]:

```
sc.tl.rank_genes_groups(adata_spatial, 'clusters', layer = 'normalized_counts',
                        method='wilcoxon', key_added = "de_genes",
                       n_genes = 30)

ranked_genes = adata_spatial.uns['de_genes']['names']
de_genes = [','.join(ranked_genes[c]) for c in ranked_genes.dtype.names]
```

### Visualize using SpatialView¶

In [ ]:

```
sample_paths = [path.join(data_path, sample_name) for sample_name in sample_names]

prepare10x_from_scanpy(adata_spatial, data_paths = sample_paths,
                       export_path = "TME",
                       cluster_genes = de_genes,
                       layer = 'normalized_counts',
                       download_repo = True,
                       launch_app = True,
                       verbose= True)
```
